# Performance of IBD machine learning classifiers varies across microbiome training data independent of geographic diversity

**DOI:** 10.64898/2026.05.21.727052

**Authors:** Giana Teresa Cirolia, Jeshua Thomas Gustafson, Anil Aswani, Ashley R. Wolf

## Abstract

Microbiome-based machine learning classifiers have strong promise for disease identification across gastrointestinal, metabolic, and immune-mediated conditions. Inflammatory bowel disease (IBD), a chronic immune-mediated disorder associated with disruption of the gut microbiome, has been a particularly successful application area. Many microbiome-based predictive models do not generalize well across independent populations and geographic contexts, despite high performance within individual datasets. We hypothesized that assembling training data by combining studies from varied geographic locations would improve model performance and generalizability. We also sought to compare the performance of different model classes across different types of training data and evaluation datasets.

We compiled seven publicly available shotgun metagenomic studies spanning five geographic regions, comprising 697 individuals with IBD or healthy controls. We trained model configurations across seven model classes and five distinct training dataset combinations and evaluated top-performing models on independent studies from the USA, Ireland, Germany, Israel, and China. Extreme gradient boosting and random forest models showed the highest and most consistent performance across training datasets.

Surprisingly, models trained on geographically diverse datasets did not outperform those trained on USA-only datasets. Instead, model performance was dependent on the evaluation study itself, with some evaluation studies consistently yielding higher performance independent of model class or training data.

Amongst models with similar AUC scores, there was a small - but consistent - overlap in the key microbial species identified. However, even for the small set of disease predictive microbes shared between models, the direction of enrichment between IBD or healthy subjects sometimes varied in opposing directions across study populations. These findings suggest that study-specific factors constrain generalization and may help explain the lack of consistent microbiome-based biomarkers for IBD.

**Importance:** Machine learning models based on the human gut microbiome are increasingly proposed as diagnostic tools for inflammatory bowel disease, but our findings suggest that identifying reliable microbiome biomarkers poses a challenge. Models trained on different datasets often selected different species as important predictors, even when diagnostic performance was similar, indicating that disease-associated microbes may depend strongly on the patient populations studied. Even species repeatedly selected across training datasets frequently showed inconsistent associations with disease, helping explain low agreement across microbiome studies. Importantly, models performed well across new patient groups independent of the geographic diversity present in the training datasets. By identifying microbial species repeatedly selected across datasets, model types, and evaluation studies, we identified a smaller group of more consistent biomarkers, including enrichment of *Klebsiella pneumoniae* and *Erysipelatoclostridium ramosum* and depletion of *Lachnospiraceae* and *Alistipes* species, which may represent stronger candidates for transferable microbiome markers.

## Introduction

Inflammatory bowel diseases (IBD), including Crohn’s disease (CD) and ulcerative colitis (UC), are chronic immune-mediated inflammatory diseases of the gastrointestinal tract that are associated with perturbations of the gut microbiome^1–6^. These disorders arise from interactions between host genetics^7^, the immune system, environmental exposures, and lifestyle factors^8–10^. Although IBD has been described for centuries, its incidence began to rise in the nineteenth and twentieth centuries in parallel with industrialization^11^. Today, IBD affects more than six million people worldwide^3^. Although the highest prevalence is still in North America and Europe, the fastest growth is now occurring across the global South in regions including South Asia, Africa, and South America^3,11–13^.

The increase in IBD cases globally starting at the turn of the 21st century, mirrors changes in the gut microbiome linked to environmental disruptions^11,14–16^. IBD risk can be modulated via microbiome-diet and microbiome-immune mediated interactions^2^. Environmental factors such as urbanization, antibiotic use, smoking, soft-drink intake, and high meat consumption are major risk factors for IBD and also linked to microbiome changes^9,12^. Conversely, exposure to animals, soil, and traditional rural environments is protective^9,17^. Migration from non-Western to Westernized environments alters the gut microbiome and increases the risk of immune-mediated diseases^10,14,18,19^. The microbiome and immune changes occur in conjunction with adoption of a low-fiber, high-fat, and highly processed Western diet^15,17,19^. Thus the global rise of IBD may be driven in part by environmentally-mediated effects on the microbiome, emphasizing the need to understand the link between microbiome disruption and IBD in different global contexts.

The gut microbiomes of IBD patients are characterized by a loss of microbial diversity, depletion of health-associated anaerobes, and expansion of pathobionts^6,10,16,20,21^. Inflammation associated with IBD alters oxygen levels, nutrient availability, and immune signals in the gut, leading to a restructuring of the microbial ecosystem^10^. These biological changes are often accompanied by reduced diversity and diminished functional gene content^22,23^. This pattern of dysbiosis is now recognized as both a marker and potential driver of disease activity, positioning the microbiome as a possible target for IBD diagnosis, prognosis, and therapy^16^. Gaps in understanding, variability in machine learning approaches, and heterogeneity across studies have hampered progress on using microbial signatures as IBD diagnostics.

As the number of IBD shotgun metagenomic datasets has grown, new opportunities have emerged to examine how IBD-associated microbiomes vary across geographic regions and populations. Shotgun metagenomic sequencing enables species level taxonomic variation, allowing precise links between microbial features and disease states^24^. However, geography and host genetics introduce nuance and heterogeneity in IBD microbiome profiles ^7,8,25^. Although many genetic risk loci are shared across populations, others are population-specific^7,26,27^. Geographic location can be as significant as disease status in explaining gut microbiome variation for many diseases, including IBD ^8,20,28,29^. Therefore, both conserved signatures of inflammation and disease, as well as region specific patterns, may define IBD profiles across countries and contexts.

Machine learning (ML) techniques are a good fit for metagenomic data since these approaches enable researchers to model data with high-dimensional, sparse, heterogeneous, and strongly nonlinear properties^30,31^. Machine learning powered studies have identified IBD-associated species including *Escherichia coli*, *Klebsiella pneumoniae*, *Ruminococcus gnavus*, *Enterococcus*, and *Fusobacterium*^6,8,22,23^. These same studies have identified putative protective species that are depleted in IBD including *Faecalibacterium prausnitzii*, *Roseburia*, *Alistipes*, and *Lachnospiraceae* species^6,8,22,23^. However, these results should be interpreted conservatively as most IBD-associated metagenomic studies focus on a single geographic area or have methodological limitations. Overfit models can result from limiting geographic diversity in training and evaluation, using supervised feature selection, or selecting models based on cross-validation training metrics. For IBD classification based on microbiome signatures, many studies train on data from single regions or limited geographic scopes and evaluate on only one or two new geographies^5,6,21,28,32^. Although recent studies have demonstrated cross-continental prediction^33,34^, many rely on pre-selected feature sets from supervised analysis prior to modeling which can inflate apparent generalizability^24,35^. For these reasons, we expand the current research in modeling IBD by extensively comparing the performance of high scoring models from training to final performance scores on seven independent and globally diverse evaluation studies.

Here, we directly tested how training dataset composition and model class influence the generalization of IBD microbiome models to new studies. We trained models on five datasets compiled from seven shotgun metagenomic studies and spanning five geographic regions. For each dataset, we trained seven model classes across regularized linear models (lasso, ridge regression, elastic net regularization), kernel-based classifiers (support vector classifier), tree-based ensemble methods (random forest, extreme gradient boosting), and neural network models to compare across complementary modeling approaches. We (i) compared cross-validation performance against true out-of-sample performance for all model classes, (ii) evaluated performance of models trained on geographically diverse versus single-region datasets, and (iii) measured how the identity of the evaluation study itself constrained accuracy. Finally, we analyzed which microbial species were consistently selected as important features across model classes and training datasets. Together, this framework enabled us to disentangle the contributions of model choice and training dataset composition on model generalization across a globally diverse set of clinical studies.

## Results

### Selection and processing of IBD metagenomic studies for meta-analysis

To compare the performance of models trained on data from United States (USA) populations versus global populations, we gathered shotgun metagenomic datasets from IBD and healthy subjects worldwide. A PubMed search yielded 950 studies that were filtered to seven eligible studies based on three criteria: availability of raw metagenomic sequencing files, accessible clinical metadata (IBD status, antibiotic use, surgery history, and age), and inclusion of both healthy and IBD patients at baseline (non-interventional) sampling. The final set included three USA studies^6,21,36,37^ and one study each from Ireland^38^, Germany^37^, Israel^37^ and China^34,39^ (Fig. 1A; Table S4).

**Fig. 1.**
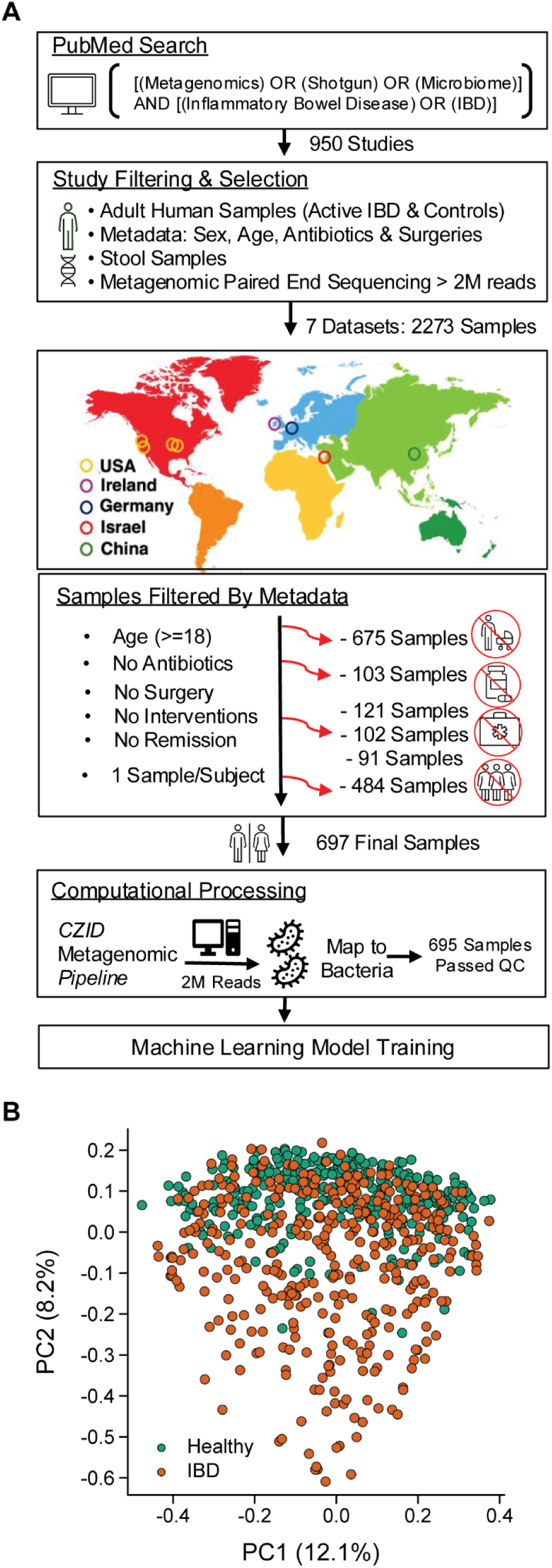
Overview of dataset selection, computational processing and patient microbiome structure. (A) A PubMed search identified metagenomic datasets from inflammatory bowel disease patients (IBD) and healthy controls. Study locations covered five unique geographies. Samples were selected based on the indicated metadata criteria and processed as raw FASTQ files through the Chan Zuckerberg CZID metagenomic nucleotide marker-gene pipeline. Bacterial species abundance was calculated as reads per million (RPM) for all samples passing quality control and the resulting species RPM count tables were used as training datasets for machine learning modeling. (B) Principal Coordinates Analysis (PCoA) of IBD patient and healthy subject gut microbiomes. PCoA based on Bray-Curtis Dissimilarity is plotted for all filtered samples (see Fig. 1A). Each point represents the gut microbiome of a single individual, IBD includes UC and CD patients.

Across these studies, samples were screened and filtered to retain adult (≥18 years) subjects with either active IBD or healthy controls. Individuals with recent antibiotic use, subjects in active clinical trials, and those with prior gastrointestinal surgery were excluded. We also retained only the first sample from each individual to reduce inter-sample correlation from studies with multiple time points. This yielded 697 samples for downstream analysis (Fig. 1A; Table S1; Table S4).We obtained raw sequencing data from the NCBI Sequence Read Archive and assigned taxonomy to each sample using the Chan Zuckerberg CZID metagenomic pipeline^40^. We retained only taxa assignments that were confidently resolved at the species level, and all mappings to genus and higher taxonomic levels were excluded from sample outputs. The resulting reads per million (RPM) count tables served as input for all modeling analyses (Fig. 1A; Methods).

Microbiome analysis of the combined studies identified differences between microbiomes from individuals with IBD compared to those from healthy controls despite differences in study geography. Principal coordinates analysis (PCoA) of these microbiome profiles identified a significant difference between IBD and healthy samples (Bray–Curtis dissimilarity, PERMANOVA, p = 1 × 10⁻⁴; Fig. 1B). Additionally, we found significantly greater dispersion among IBD samples compared to healthy controls (PERMDISP, p = 1 × 10⁻⁴), indicating increased inter-individual heterogeneity in disease-associated microbiomes. When tested individually, all but one dataset showed significant separation between IBD and healthy samples (PERMANOVA p < 0.01; Table S2; Fig. S1). Across all datasets, PCoA still revealed clustering by study populations (Fig. S8). Interestingly, ecological separation was not organized strictly by geography, as all studies (including the 3 USA study populations) showed significant separation from one another (Fig S8).These findings suggest that study-associated microbiome differences were not determined solely by geographic origin, but also reflected more nuanced study-intrinsic ecological and technical factors.

### Models trained on different training data exhibited stable and class-specific cross-validation scores

We trained models across five distinct training datasets constructed by combining three independent studies (Table 1, Table S1, Table S4). The training datasets were selected to test how including different amounts of geographic diversity in training affected model behavior. One training dataset included three U.S. studies (labeled USA), while four training datasets used combinations of three international studies (labeled Int_1 through Int_4) (Table 1; Table S1). For each training dataset, we trained the same seven model classes—random forest (RF), extreme gradient boosting (XGBoost/XGB), lasso, ridge regression (ridge), elastic net regularization (ENet), neural network (NN), and support vector classifier (SVC).—across a broad hyperparameter space (see Methods). This yielded an average of ∼7,000 models per class per training dataset (Table S3) and a total of ∼249,000 models across all model classes and training datasets. Because training datasets contain distinct patient compositions, this design enabled us to assess each model class’s sensitivity to changes in underlying data distribution.

**Table 1.** Composition and metadata characteristics of training data. Training datasets include three studies which balance (i) IBD to Healthy ratio and (ii) total subjects per training dataset. Studies named by country and last author or the name of the research dataset. International abbreviated as Int_1.

| Training Datasets | Studies | Male | Female | Age (Median & IQR) | IBD | Healthy | Total |
| --- | --- | --- | --- | --- | --- | --- | --- |
| USA | USA_Elinav, USA_HMBP & USA_PRISM | 84 | 97 | 36 (28–54) | 220 | 99 | 319 |
| Int_1 | China_Jia, Germany_Elinav & Israel_Elinav | 162 | 144 | 44 (30–58) | 137 | 180 | 317 |
| Int_2 | China_Jia, Germany_Elinav & Ireland_Hill | 158 | 147 | 44 (31–58) | 118 | 195 | 313 |
| Int_3 | Germany_Elinav, Ireland_Hill & Israel_Elinav | 127 | 162 | 50 (37–60) | 112 | 196 | 308 |
| Int_4 | China_Jia, Ireland_Hill & Israel_Elinav | 108 | 69 | 31 (24–42) | 95 | 101 | 196 |

For the thousands of model configurations in our training results, we calculated both the average 5-fold CV score and its dispersion for each of the 5 training datasets, separated by model class. We hypothesized that, per model class, the average 5-fold CV scores across the five training datasets would serve as a proxy future generalizability. We expected that model classes with higher and tightly clustered CV scores would have consistent and relatively stronger scores when evaluated outside of training. Our analysis showed that mean 5-fold CV performance (the average AUC across all five folds of training) remained largely consistent within each model class regardless of training dataset (Fig. 2). XGB achieved the highest and most stable performance (mean CV range = 0.863–0.898; SD range = 0.007–0.017), followed by RF models (mean CV range = 0.853–0.874; SD range = 0.018–0.055) (Fig. 2). Comparatively, ridge and neural net had intermediate performance, while ENet, lasso, and SVC had the lowest mean performance and highest variability (Fig. 2). Most model classes maintained their class-specific 5-fold CV performance averages across all training datasetsHowever a subset of model classes (SVC, ENet, lasso) were more sensitive to differences in training datasets - having both lower average performance and more performance variability (Fig. 2). We next tested whether the mean 5-fold CV and CV variability for a given model class would predict performance trends on out-of-sample evaluation studies.

**Fig. 2.**
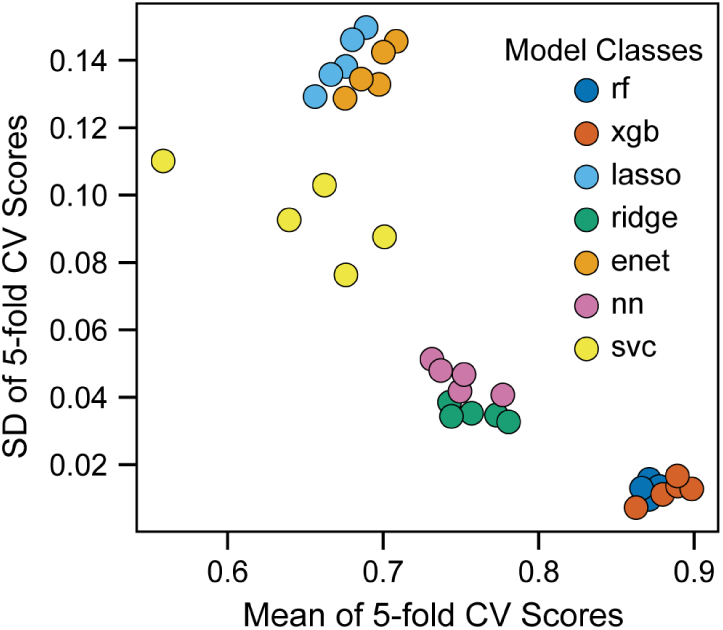
Mean cross-validation performance versus variability within each training dataset. Each colored point represents the average 5-fold cross validation (CV) (x-axis) and standard deviation (SD) (y-axis) calculated across all models trained in a training dataset. Average and standard deviation are shown for the 5 training datasets (see Table 1) separated per model class.

### Differences in model class performance are less pronounced on out of sample evaluation

To evaluate how training 5-fold-CV scores translates to real-world generalization, we tested each top scoring model on four fully independent evaluation studies from diverse geographic locations (Fig. 3A). We selected the top 13 scoring models (ranked by 5-fold CV score) per model class in each training dataset and tested their performance on four withheld studies (455 total models, 1,820 total out of sample AUCs; Fig. 3A). Specifically, models trained on the USA dataset were evaluated on studies from Ireland, Germany, Israel and China, while models trained on the international datasets were evaluated on these locations, as well as 3 studies from the USA (Fig. 3A). This setup enabled us to explore how model classes perform when tested on many clinical studies with variation in demographics, geography and batch effects/study design.

**Fig. 3.**
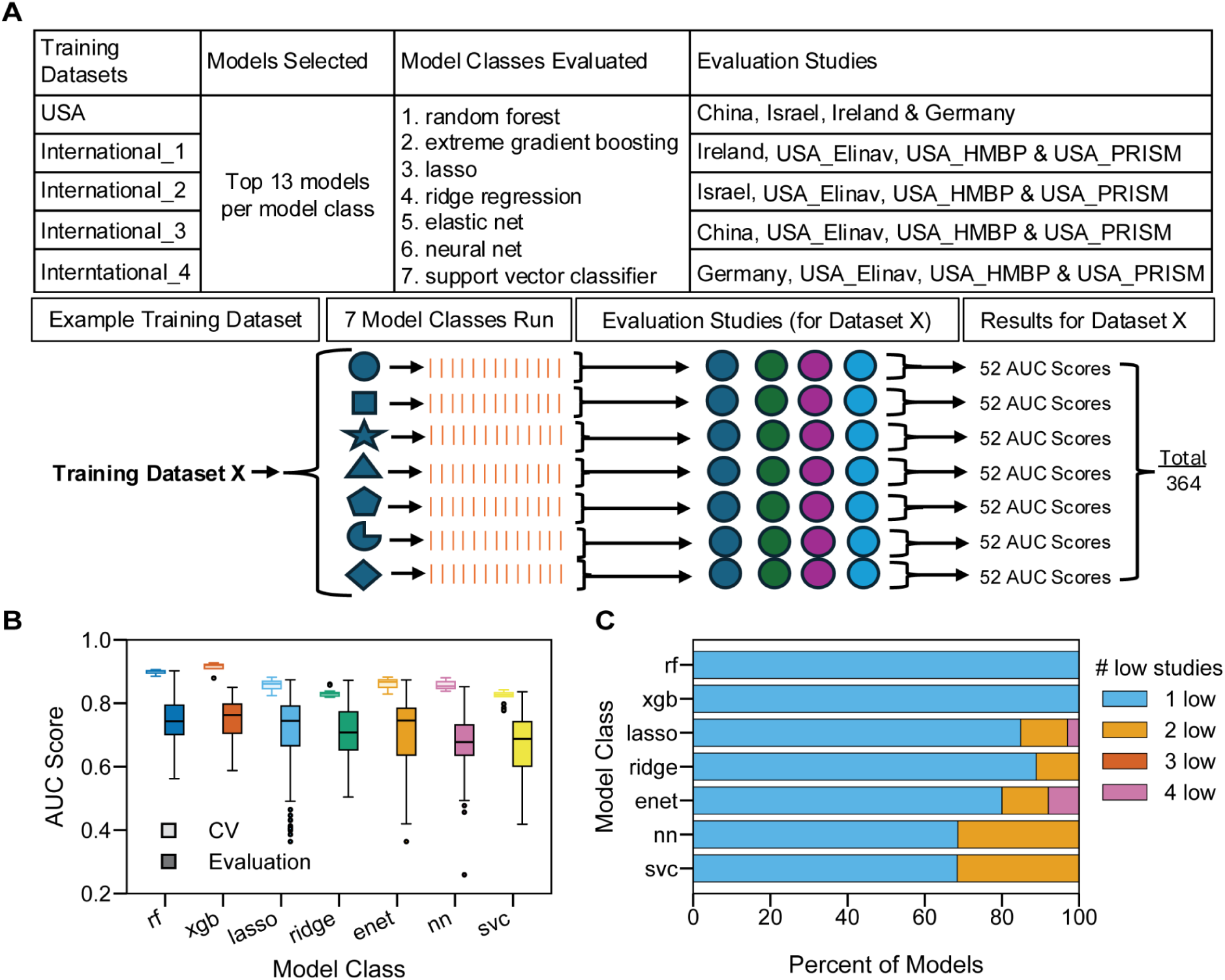
Performance comparisons of selected top models in training and on evaluation studies. (A) For each training dataset, the top 13 models per model class were evaluated on withheld studies. Schematic shows the evaluation process for an example training dataset. (B) Boxplots compare 5-fold cross-validation (CV) AUC scores from training (color outlined boxes) with all out-of-sample evaluation AUC scores (color filled boxes) for all top models per class. Center lines indicate medians, boxes represent interquartile ranges, and whiskers extend to 1.5× the interquartile range. (C) Stacked horizontal bars show the percentage of top models per class with 0–4 low-scoring evaluation studies.

The results for 5-fold CV when using only the top 13 models was consistent with the findings above. 5-fold CV scores for the top 13 models were tightly distributed, with model-level 5-fold CV medians ranging from 0.83–0.92 (IQRs 0.008–0.024) (Fig. 3B, outlined boxes). Extreme boosting gradient and RF maintained their advantage relative to other model classes, though the median performance differences were minimized when analysis was limited to the 13 high performing models. In contrast, out-of-sample evaluation AUCs exhibited broader dispersion (Fig. 3B, color filled boxes), indicating more performance variability between model classes. XGB (median = 0.76, IQR = 0.09) and RF (median = 0.74, IQR = 0.09) showed the highest median evaluation AUCs and the smallest dispersion of scores. Notably ridge, lasso, and ENet had lower median performance (0.71–0.75) with wider interquartile ranges (0.12–0.15). Neural networks and SVC were the lowest median evaluation AUCs (0.68-0.69) (Fig. 3B). These performances generally reflect their ordering in 5-fold CV scoring as well (Fig. 3B). Thus, the model class differences observed in training are maintained albeit attenuated when evaluated on withheld studies.

To further assess differences in stability across model classes we counted how often models within a class fell below a set threshold AUC score. We hypothesized that model classes with more generalizability would have fewer models that scored low on any of their evaluation studies. We defined the threshold of low performance as AUC < 0.61 (one standard deviation below the mean of all 1,820 evaluation AUCs) (Fig. 3C). Approximately 80% of RF and XGB models showed no low-performing studies (Fig. 3C, green bars), whereas SVC models frequently had low-performing studies (Fig. 3C, orange, purple & pink bars). Other model classes showed intermediate behavior, but still more poor evaluation AUCs compared to XGB and rf. This result suggests that XGB and RF may have the most predictable generalizability to new studies.

### Model class predictions have low correlation and ensembling classes does not improve performance

We next asked if model classes make similar predictions on the same individual patients. Model classes have overlapping AUC performance distributions and we wanted to know if they categorized individuals similarly. We hypothesized that models can score similarly while classifying patients differently. This disagreement may reflect complementary signals in the microbiome which might improve AUC scores if model classes were combined. To test this, we focused our analysis on single model classes trained on the USA dataset (Table. 1, USA dataset composition). For each model class, we averaged predictions for the top models across four studies (Ireland, Germany, Israel & China) and computed pairwise correlations between the model classes (Fig. 4A). Correlations between model classes varied widely ranging from .22 to 93 (Fig 4A). The most correlated models were ENet with lasso (r = 0.93), and RF with XGB (r = 0.80). Neural networks differed the most from other classes, with correlations ranging from r = 0.22 to 0.36. These patterns suggest that underneath similar AUC scores, model families classify patients differently.

**Fig. 4.**
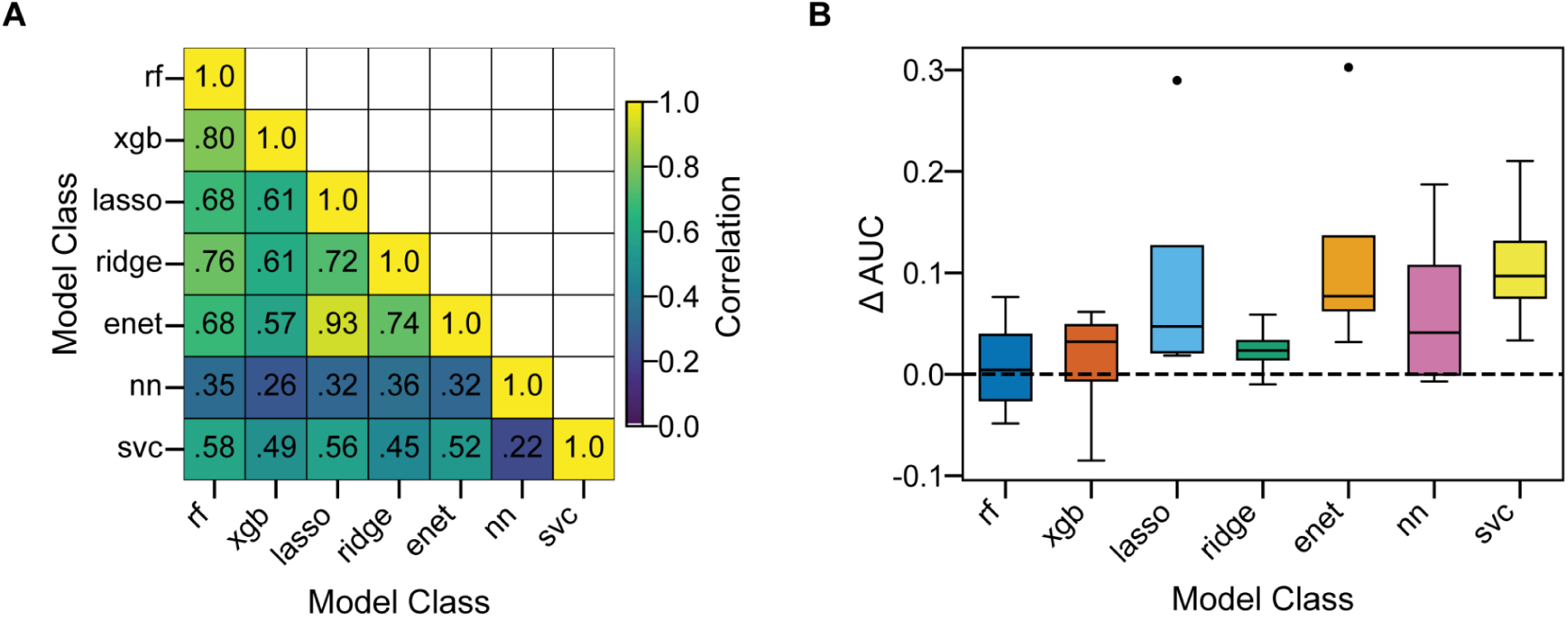
Correlation of model classes and comparisons of ensemble strategies. (A) Correlation of model class ensembles for the USA training dataset. Predictions are pooled across evaluation studies (Ireland, Germany, Israel & China). Predictions are averaged across the top 13 models per class. Pairwise Pearson correlations between ensemble predictions are shown. (B) 7-class versus single class ensemble scores for models from the USA training dataset. Ensemble models are evaluated on four international studies (Ireland, Germany, Israel & China). Per class, single model class ensembles are compared to seven-model class ensembles per study. Box plots show the AUC difference per study (ΔAUC = AUC (single model ensemble) − AUC (7 model class ensemble)). Center lines indicate medians, boxes represent interquartile ranges, and whiskers extend to 1.5× the interquartile range.

Next we tested whether ensembling models of different classes could improve performance. We hypothesized that if model classes rely on different features to classify patients, combining model classes may improve AUC scores. To test this we compared single model class ensembles (average predictions of top 13 models per class) to the ensembles of all seven model classes (combining the 13-model ensembles together across all seven classes). We evaluated these two ensembling strategies on the four international studies (Fig. 3A, evaluation studies). To measure differences we calculated the AUC score difference between the single-class and seven-class ensembles for each study (Fig. 4B). The seven class ensemble performed generally better than weaker models (Fig. 4B), but did not out perform the higher scoring individual model classes like RF and XGB (Fig. 4B). Overall, seven-class ensembles raise and stabilize performance of weaker model families but do not improve performance for models like XGB and RF (which already perform well on their own).

### Model performance is not improved by adding geographic diversity to the training data

Next, we tested whether the geographic diversity of a training dataset affects a model’s ability to perform well on several withheld global studies. We hypothesized that models trained on pooled international datasets would outperform models trained on a single country when tested on datasets from Ireland, Germany, Israel and China (Fig. 5A). We performed head to head comparisons for all 7 model classes and all 4 evaluation studies. We saw that both international-trained models and USA-trained models scored higher than their comparison pair a roughly equal number of times (Fig. 5B-E, international-trained models have higher AUCs in 15 of 28 comparisons (53.6%), USA-trained models have higher AUCs in 13 of 28 comparisons (46.4%.)). Additionally for all comparisons between models trained on USA or international datasets, the difference in AUC scores were not significant (Paired Delong Test, p > .05). Still, a slight dataset-specific substructure emerged (Fig. 5B-E). We saw that USA-trained models generally performed better on China (6/7 ensembles) and Germany (5/7 ensembles), whereas international-trained models performed better on Israel (6/7 ensembles), with Ireland showing near parity (Fig. 5B). Thus overall, the one training regime (international versus USA) did not produce generally better performing models on all evaluation studies, but often specific training datasets slightly but consistently improved performance for individual studies.

**Fig. 5.**
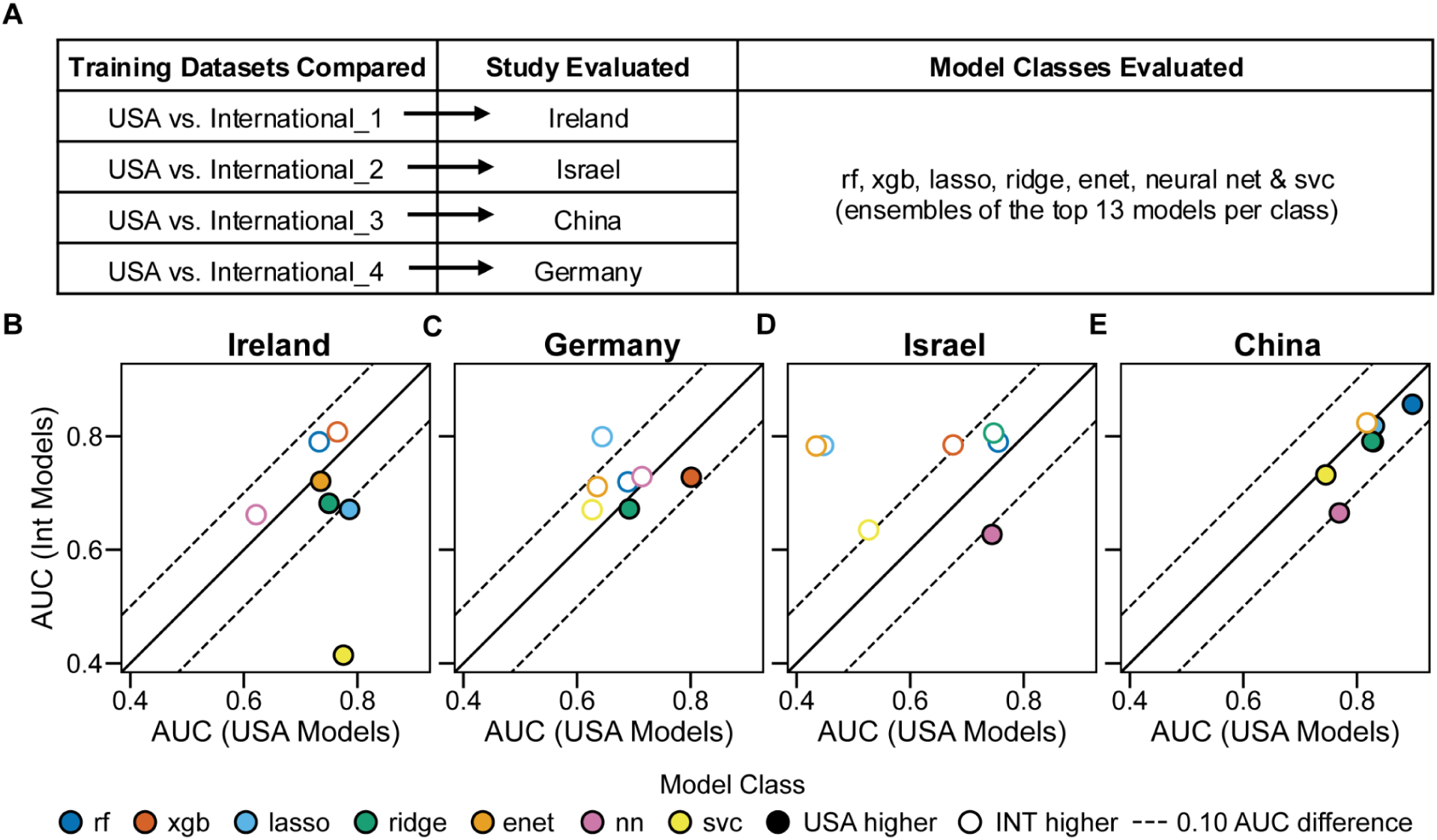
Performance comparison for models trained on USA versus international training datasets. (A) Schematic of USA versus International training dataset comparisons. AUC scores from models trained on the USA dataset are compared with models trained on international datasets (Int_1 to Int_4). Paired comparisons are made on 4 international studies separated by model class (Ireland, Germany, Israel & China). (B-E) For each international evaluation study (Ireland, Germany, Israel & China), ensemble AUC scores from the USA training datasets are plotted (x-axis) against ensemble AUC scores from the paired International training dataset (y-axis). Each point represents a model class ensemble. Dashed lines indicate AUC differences of 0.1.

### Models trained on different international datasets perform comparably across USA evaluation studies

We next compared models trained on two international datasets by evaluating them on three USA studies (Fig. 6A). Here we saw a similar trend to our previous tests (Fig. 5B-E). Models trained on one international dataset scored higher than models trained on the other international dataset roughly an equal number of times across all comparisons. As before, these training datasets also exhibited small study-specific preferences. Models trained on dataset Int_3 consistently outperformed those from dataset Int_4 on USA_HMBP (6/7 ensembles), while models from dataset Int_4 outperformed those from dataset Int_3 on USA_PRISM (7/7) (Fig. 6B-D). Still, no differences reached significance (Paired Delong Test, p-value >.05). Thus it appears that internationally trained models perform comparably well on a variety of USA studies irrespective of input training data. Furthermore, the small biases for better performance on specific evaluation studies continued to occur even when models were trained on highly similar datasets (Table. 1, ⅔ shared input data between international training datasets). These biases were equal magnitude to those seen previously when we compared models from the USA dataset versus international datasets, where no input studies were shared between training datasets (Table. 1, training data composition). These results indicate that even a small change in training dataset composition can create a small performance bias for specific evaluation studies.

**Fig. 6.**
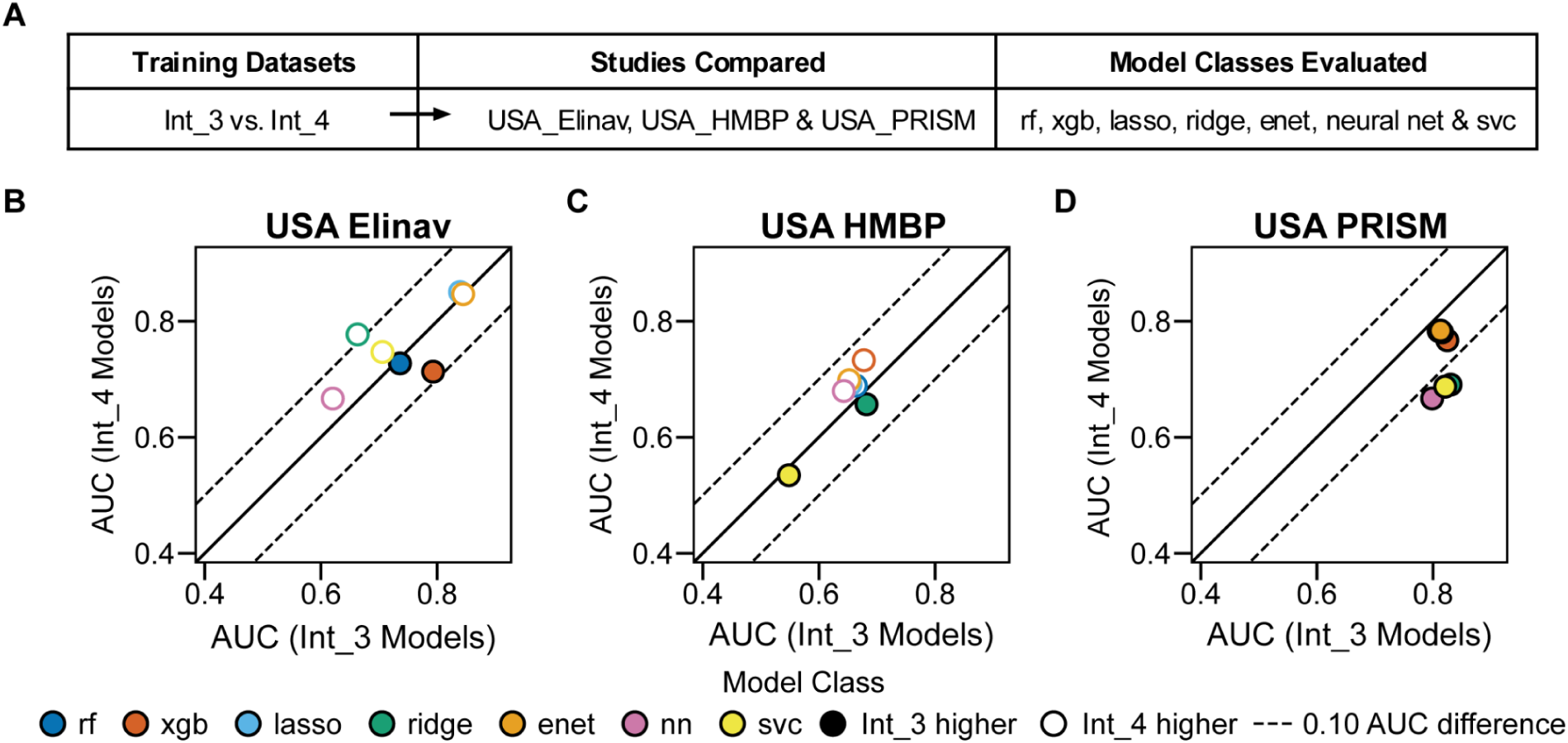
Comparison of models trained on international training datasets when evaluated on USA studies. (A) Schematic of model performance comparisons for international training datasets. Comparison of AUC scores for models from two international training datasets (Int_3 vs. Int_4) when evaluated on 3 USA studies. Top 13 models (by CV score) ensembled per model class. (B-D) Comparison of single–model-class AUC scores from Int_3 versus Int_4 training datasets on USA evaluation studies. Scores from models from Int_3 are plotted (x-axis) compared with paired counterparts from Int_4 (y-axis). Comparisons made for 3 USA studies. Each point represents one single model class ensemble (13 top models per class) evaluated on the same study. Dashed lines indicate AUC differences of 0.1.

### Performance differences are driven by differences in evaluation studies

We next tested whether the evaluation studies constrain model performance, independent of the effect of model class or training dataset. Given the modest difference in performance between models training on different training datasets, we hypothesized that achievable AUCs per evaluation are influenced by the properties of each evaluation study - perhaps more strongly than by differences between model class or training dataset. To test this, we computed out-of-sample AUCs for all single–model-class ensembles (top 13 models per model class) across all applicable datasets and summarized this performance in box plots distributions (Fig. 7A, international studies; Fig.7B, USA studies).

**Fig. 7.**
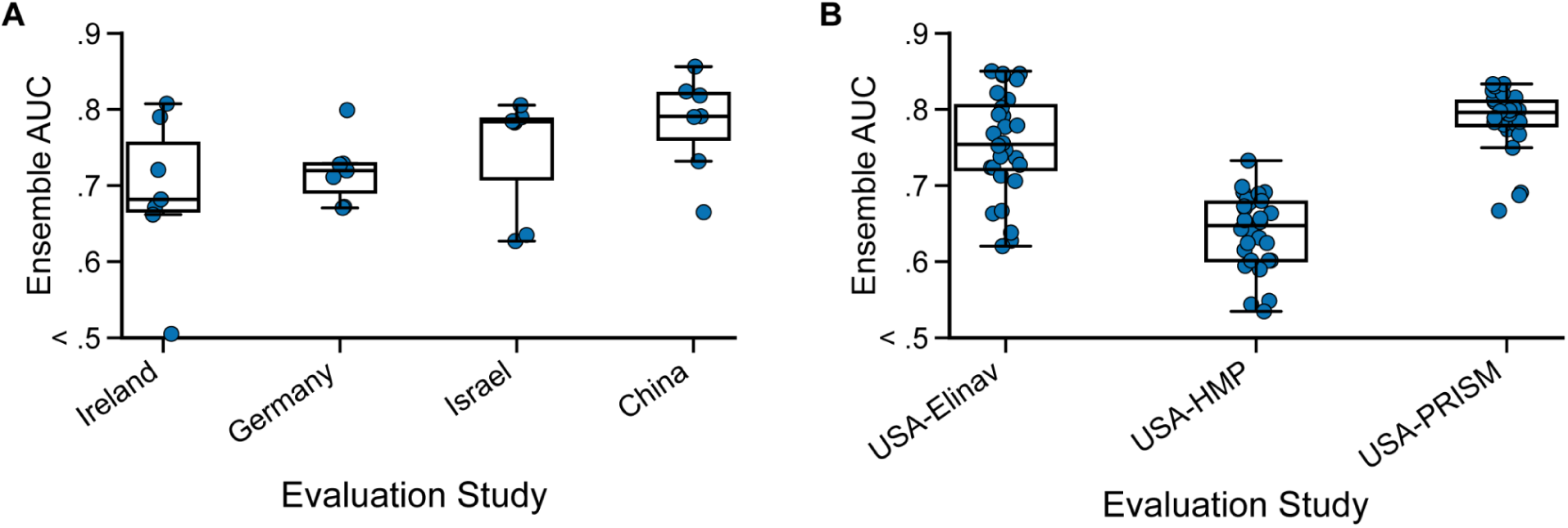
General AUC performance scores across all evaluation studies. (A) Boxplots show the distribution of AUCs for international evaluation study (single–model-class ensembles, studies: Ireland, Germany, Israel & China) across all applicable training datasets (USA and Int 1-4). Each point represents one ensemble AUC score.. Boxes indicate the median and interquartile range (IQR), with whiskers extending to 1.5 IQR. (B) Boxplots show the distribution of AUCS on USA evaluation studies (single–model-class ensembles, studies: USA_Elinav, USA_HMBP & USA_PRISM) across all international training datasets (Int_1–Int_4). Each point represents one ensemble AUC score. Boxes indicate the median and interquartile range (IQR), with whiskers extending to 1.5 IQR. Y axis trimmed for visualization.

Across the four international studies (Ireland, Germany, Israel & China), we observed median differences in achievable performances regardless of the model classes or training datasets (Fig. 7A). When aggregating across all model classes, median AUCs ranged from 0.67 (IQR 0.05) for Germany to 0.79 (IQR 0.06) for China (Fig. 7A). These median differences persisted across all four international studies (relative ranking China, Ireland, Israel & Germany), indicating that evaluation study-specific factors constrain performance.

To assess whether these dataset-level differences persist even among the strongest and most stable model classes, we repeated the analysis but restricted ensembles to XGB and RF (Fig. S2). As expected, for XGB and RF, the median AUCs increased for all datasets and within-dataset variability decreased (Fig. S2). However, while the performance rose for all datasets, the relative ordering of datasets remained unchanged: China consistently showed the highest performance, while Germany remained the lowest (Fig. S2). Thus, stronger model classes improved AUC scores for all studies but did not decrease the separation between median performances across studies.

We hypothesized that these differences were not due to geographic difference between the studies but due to the heterogeneity of microbiome samples between studies regardless of geography. To test this hypothesis, we focused on 3 datasets from the same country (USA_Elinav, USA_HMBP & USA_PRISM) and plotted the AUC performance metrics across all models and training datasets. A similar pattern to the international studies was observed across the three USA cohorts. Across all international training datasets (Int_1 – Int_4) and all 7 model classes, median AUCs ranged from 0.63 (IQR 0.06) for USA_HMBP to 0.77 (IQR 0.05) for USA_PRISM (Fig. 7B). The differences were consistent across individual training datasets, suggesting that properties of each USA study impact achievable performance (Fig. S4). These patterns were also maintained even when the strongest model classes (XGB and rf) were evaluated separately (Fig. S3).Together, these results demonstrate that IBD studies may differ in how difficult they are to classify accurately. The upper bound on achievable AUC appears to be largely set by evaluation study-specific factors, with model choice and training composition modulating performance within these constraints.

### There is a limited set of shared “important features” (species) across model classes and training datasets

We next extracted features with the top importance scores from the highest scoring five models (by 5-fold CV) per model class across all five training datasets (Fig. 8A, 125 models total). We sought to determine whether top features were shared or distinct across model classes and training datasets. To standardize the number of features selected per model, we used the lowest number of features across all model classes (80 features) needed to capture 95% of total feature importance (Fig. S5). For each model class and each training dataset - these 80 features were consolidated into a unique set across the top 5 models (Fig. 8A). Pooling the top-80 features across per model class and training dataset yielded ∼90–160 unique taxa per class, far fewer than the theoretical maximum of 400 (80 features x 5 top models) (Fig. S6). This consolidation indicates agreement within individual model classes on key features. Across training datasets, only ∼70 taxa appeared in all five training datasets (Fig. S7), showing that training composition strongly influences which features models emphasized. Nevertheless, this shared set represents a potentially stable microbiome signal across geographies. We ranked these shared taxa by total frequency across all models (5 top models x 5 models classes x 5 training datasets = 125 possible occurances) and focused on the top 25 most occuring species (Fig. 8B). This “global core” included many organisms previously linked to IBD, including *Escherichia coli* (107/125 models), *Erysipelatoclostridium ramosum* (97/125), *Lachnospiraceae bacterium sunii* (87/125), *Alistipes communis* (85/125), *Eggerthella lenta* (82/125), *Klebsiella pneumoniae* (80/125), and *Butyricimonas faecalis* (77/125), along with additional *Alistipes*, *Bacteroides*, and *Lachnospiraceae* species.

**Fig. 8.**
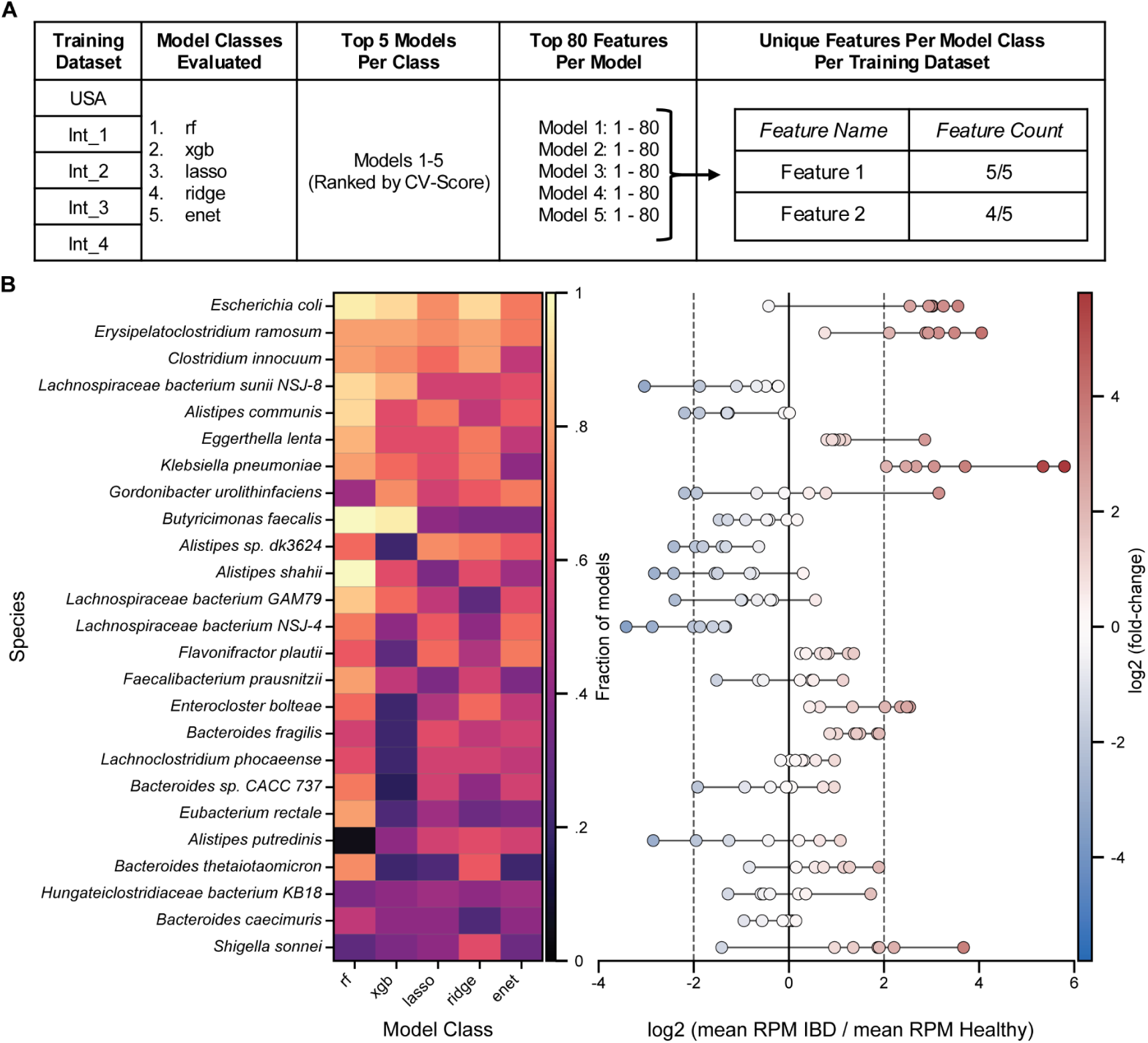
Selection of important features (species) per model and distribution of top species across studies. (A) Schematic of feature selection per model. The top 80 features were extracted from high performing models (top 5 by CV score). For each model class and training dataset combination, the non-redundant features were retained as well as the count of occurrences (1/5 = min occurrences, 5/5 = max occurrences). (B) Heatmap showing the frequency for top occurring microbial species across all model classes and training datasets. Heatmap is restricted to the top 25 most recurrent species by total counts. Rows correspond to species, ordered by total count of occurrence per species across all models and datasets. Columns correspond to model classes. Heatmap values range from 0 to 1 (out of 25 possible models - 5 model classes x 5 top species). (C) Log₂ fold-change of mean species abundance (IBD patients/ Healthy subjects) per study for the top 25 taxa. Points correspond to Log₂ fold-change (IBD mean/ Healthy mean) per study (Ireland, Germany, Israel, China, USA_Elinav, USA_HMBP & USA_PRISM). Red indicates higher abundance in IBD patients, and blue indicates higher abundance in Healthy controls.

Some of these “global core” taxa show consistent enrichment or depletion signatures across all studies while others have opposing trends between data sets. We computed the mean abundances of the “global core” taxa in IBD and Healthy subjects (RPM) for each study and summarized enrichment as the log₂ fold-change (IBD / Healthy) (Fig. 8C, see methods). Several taxa were consistently depleted in IBD across all studies, including *Lachnospiraceae bacterium NSJ-4*, *L. sunii NSJ-8*, *Oscillibacter sp. NSJ-62*, *Coprococcus sp. ART55/1*, and *Alistipes sp. dk3624*. In contrast, *Klebsiella pneumoniae* and *Erysipelatoclostridium ramosum* were consistently enriched in IBD across all datasets. However, many taxa showed intermediate (no large fold change) or study-dependent behavior, with enrichment direction varying by dataset (Fig. 8C). Thus, even when models converge on the same taxa, the strength and direction of their association with IBD patients varies across datasets.

## Discussion

In this study, we investigated how machine learning model class and training dataset composition influence the cross-study generalizability of microbiome-based classifiers for inflammatory bowel disease (IBD) across geographic contexts. Developing microbiome models that generalize robustly across populations remains a major challenge because microbiome composition itself is strongly shaped by geography, migration history, industrialization, diet, lifestyle, and environmental exposure^8,14,18,41^. In line with this, previous studies have demonstrated substantial microbiome restructuring associated with westernization, migration, and dietary transition^14,18^, while intercontinental analyses of IBD patient populations have shown that geography can contribute substantially to microbiome variability between studies^8,34^. Together, these findings raise important concerns regarding whether microbiome-based diagnostic models trained in one population can reliably transfer across independent geographic study populations.

Many microbiome machine learning studies report strong predictive performance within individual studies and, increasingly, across external validation datasets^28,30^. However, performance often declines during evaluation on independent studies, highlighting the difficulty of developing microbiome classifiers that generalize robustly across populations and study designs^8,28,30^. These challenges are particularly relevant in microbiome research where disease status may explain only a modest fraction of total microbiome variation^8,34^ and where technical, environmental, dietary, and host-associated factors can differ substantially between study groups^6,32^. Together, these observations suggest that microbiome classifier performance may depend not only on disease-associated biology, but also on broader study-specific ecological and technical context.

Given these challenges, we hypothesized that increasing geographic diversity within the training data would improve model transferability by exposing models to a broader range of microbiome states and encouraging learning of more generalizable disease-associated patterns. We also hypothesized that testing models on multiple withheld evaluation studies of the same disease (IBD), would provide evidence of whether models can achieve stable generalization. Surprisingly, increasing geographic diversity in the training datasets did not consistently improve predictive performance. Models trained on pooled international studies performed comparably to models trained on pooled U.S.-only studies despite the expectation that broader microbiome diversity would strengthen transferability. Furthermore, predictive performance appeared to be driven primarily by the evaluation dataset itself, with some studies consistently yielding higher or lower classification accuracy regardless of training dataset composition or machine learning architecture. This pattern suggests that certain studies may intrinsically contain a stronger or weaker disease-associated microbiome signal. Studies with stronger inflammatory-associated microbial restructuring^5,6,29^ may therefore yield clearer classification boundaries and improved predictive performance, while technical variation, past medication exposure, environmental factors or diet may further contribute to study-specific signal strength. Together, these findings suggest that simply increasing geographic diversity in training data may not be sufficient to overcome the broader sources of variability that exist between microbiome studies.

Although predictive performance remained relatively stable across model classes, different models and training datasets frequently selected distinct microbial features. This highlights an important issue in microbiome machine learning: strong predictive performance does not necessarily imply stable biological interpretation. Different models may achieve similar classification accuracy while relying on a largely non-overlapping set of microbial features, particularly in high-dimensional microbiome datasets where many correlated taxa contain similar ecological information^30,32,35^. Prior studies have similarly demonstrated that different feature selection approaches can identify inconsistent microbial biomarkers in IBD datasets^31^, highlighting the instability that can emerge during microbiome feature selection and biomarker interpretation^30,35^. At the same time, our modeling approach allowed us to identify a subset of microbial taxa that repeatedly emerged across multiple model classes and training datasets. These taxa suggest the presence of a more stable underlying inflammatory-associated microbiome signal. However, these recurrent taxa should still be interpreted cautiously, as many organisms repeatedly associated with IBD are also implicated across broader inflammatory and dysbiotic states rather than uniquely disease-specific processes^5,6,28,29^. Cross-disease microbiome analyses have similarly suggested that some microbial features associated with IBD may represent more general inflammatory ecological signatures rather than uniquely IBD-specific biomarkers^28,30,42^.

Importantly, many of the consistently selected taxa in our analyses overlap with previously reported hallmarks of IBD dysbiosis, including enrichment of *Enterobacteriaceae*, *Escherichia coli*, and *Klebsiella* pneumoniae alongside depletion of protective taxa such as *Faecalibacterium prausnitzii*, *Roseburia*, *Alistipes*, and members of *Lachnospiraceae*^5,6,20,21,29^. Prior studies have suggested that inflammatory-associated organisms such as *Enterobacteriaceae* and *Escherichia coli* may expand under oxidative and inflammatory gut conditions^5,6,21^, while many depleted taxa are linked to butyrate production, epithelial maintenance, and immune regulation^5,6,29^. Together, these findings suggest that while individual microbial species may vary substantially across studies and populations, broader ecological shifts between inflammatory-associated and protective microbial communities remain one of the clearest and most reproducible signatures of IBD. These observations further suggest that microbiome-based diagnostics and therapies may ultimately benefit more from focusing on microbial genes, metabolic pathways, ecological interactions, or inflammatory community structure rather than relying exclusively on individual taxa as biomarkers^5,6,24,29,32^.

Our study is limited by the publicly available datasets currently accessible for large-scale metagenomic comparison. These studies differed in sequencing protocols, sample handling, preprocessing pipelines, and metadata completeness. They are also limited in geographic range, with major underrepresented or missing global regions in our datasets include Africa, South Asia, Southeast Asia, Latin America, Oceania, and Eastern Europe/Central Asia due to limited shotgun metagenomic IBD research studies in those areas. Although all samples were reprocessed through a unified analytical workflow and filtered using consistent criteria, residual technical variation may have contributed to study-specific performance effects and represents a challenge that would also likely affect real-world clinical deployment of microbiome-based diagnostics. Additionally, metadata related to diet, medication use, environmental exposures, and disease severity were not consistently available across studies and may contribute substantially to microbiome variation and classifier performance. Furthermore, to increase sample size and maintain statistical power across datasets, ulcerative colitis and Crohn’s disease were grouped into a single IBD class despite their known biological heterogeneity, although similar combined disease frameworks have been used in previous microbiome modeling studies^29^. Within this experimental set up, the fact that all model classes and training data compositions provided reasonably performing models, indicates that allowing for a degree of technical and biological variation in the training data may actually bolster models against overfitting to singular or highly cleaned training sources.

Overall, our findings highlight both the promise and the limitations of microbiome-based machine learning models for IBD. While predictive models can achieve strong performance within certain studies, developing classifiers that generalize consistently across populations and studies remains difficult due to substantial microbiome heterogeneity between datasets. At the same time, the repeated emergence of shared inflammatory-associated ecological patterns across models and studies suggests that there may still exist broadly conserved microbiome signatures of inflammatory disease. Future progress will likely require larger international datasets and greater emphasis on microbial function and ecology in order to develop more robust and globally transferable microbiome-based diagnostics.

## Data Availability

## Data and Metadata

All processed patient metadata and metagenomic abundance data generated in this study are stored in a SQLite database (bio.sqlite3). The database includes species-level abundance tables, associated metadata, and model outputs. The full database and accompanying schema (.schema) will be made publicly available upon publication and are available upon request.

All sample original SRA accessions and study origins located in Table S4.

## Code Availability

All code used for data processing, model training, and analysis is publicly available at: https://github.com/Jtth3Brick/bio-sklearn-trainer

https://github.com/Jtth3Brick/bio-experiment

## Acknowledgments

This work was supported by the National Institutes of Health (NIH) grant R35 GM147513 and the associated Diversity Supplement, as well as the National Science Foundation (NSF) Digital Transformations of Development (DTOD) Fellowship.

## Author Contributions

G.C., A.A., A.W., and J.G. conceived and designed the computational analyses.

G.C. and J.G. analyzed the data.

G.C., A.A., A.W., and J.G. wrote the manuscript.

J.G. implemented and tested the machine learning and modeling code.

All authors reviewed and approved the final manuscript.

## Materials and Methods

### Public Datasets: Study Search And Selection Criteria For IBD and Controls

A PubMed search was performed using the key words (((metagenomics) OR (shotgun)) OR (microbiome)) AND (Inflammatory Bowel Disease) OR (IBD)) for which 950 studies were searched/returned (additionally studies cited by these 950 were also included if they met the below criteria search)

From these queries, studies were screened to ensure datasets were:

1. Shotgun metagenomic data from IBD patents and/or controls
2. 150-250 base pair paired-end Illumina short read sequences
3. Average of =>2M reads per sample per study
4. Included baseline samples if interventional studies
5. Accompanied by subject metadata for age, antibiotic use, history of gastrointestinal surgeries, and IBD disease status index (active IBD or remission as determined by individual criteria set per study). Note: Occasionally studies had missing data for specific age - patients with missing age were still included if the study noted that all patients were over 18.
6. All studies required to have both IBD and healthy subjects collected per datasets.

Our screening resulted in 8 data sets (clinical groups of patients and controls). Of these 8 datasets (1 set) combined 2 studies using patients that were recruited and treated in the same patient medical centers (same local patient population, and data collection/processing pipelines but different subjects). As these samples were processed and sequenced in the same experimental labs, we combined samples from the same medical center into one “dataset” per medical center. The remaining studies were kept as individual study datasets leading to a total of 7 datasets used in our modeling. Of these 7 studies, 3 = USA, 2 = Europe (Ireland, Germany), 1 = Middle East (Israel) and 1 = Asia (China). Across these studies there were 2,273 metagenomic samples (Table S4, SRA ascension numbers)

## Data storage and database structure

All data used in this study were stored in a single SQLite relational database (bio.sqlite3), which integrates metagenomic sequencing outputs, patient metadata, machine-learning model training results and machine-learning model evaluation results. This database serves as the sole source of input for all analyses and figures presented in this manuscript. The database contains three main classes of information:

1. metagenomic abundance data (rpm)
2. patient metadata
3. machine-learning training results
4. machine-learning evaluation/prediction results
5. extracted feature importances for top models
6. “describe item 6”

Within this database there is a built in .schema file overviewing all contents. =SQL queries, data filtering logic, and analysis workflows are implemented in the accompanying Jupyter notebooks. The complete database schema is provided within the database itself as a .schema component of the database that can be accessed through SQL queries.

## Filtering Of Metadata Included For Studies

To harmonize metadata across samples (and for machine learning analysis) we filtered our input 2,273 samples to only samples with metadata available for the following fields: age, condition (UC, CD, healthy), remission, intervention, surgery.

We applied a filtering procedure using SQL logic based on the following criteria:

1. Age 18 or over (age >=10) - either noted in metadata as individual ages or in study text as >=18 for all.
2. Not following any interventional diet modification at the time of sampling (intervention = 0)
3. No antibiotic treatment recorded within three months of consent (antibiotics = 0)
4. No patients with history of gastrointestinal surgery or resections (surgery = 0)
5. No patients in remission or inactive disease status (as determined by the study criteria) (remission = 0)

Note* Patients currently receiving medical therapy for their baseline treatment are still included if that therapy is part of their medical care however,

7. Samples taken during interventions for patients enrolled in a current interventional trial were not included (intervention = 0), but patients who previously participated in interventional studies were included at other times where.
8. Only one sample (the first sample recorded) per patient is taken for longitudinal studies with multiple timepoints (patient_id = unique identifier per subject and run=sample per subject). This equates to a baseline sample for all patients.

### Shotgun Metagenomic Sequence Processing and Model Input Data

All raw short-read fastq sequence files (from human stool metagenomic samples) for each selected study were downloaded from SRA and processed through the Chan Zuckerberg CZID metagenomics pipeline Version 8 to ensure consistency in metagenomic mappings (See table S4 for SRA ids).

In short, the CZID pipeline uses the following steps:

1. Host filtration and removal of ERCC sequences using STAR
2. Trim sequencing adapters with Trimmomatic
3. Quality filter using PriceSeq
4. Identify duplicate reads using czid-dedup
5. Filter out low complexity sequences using LZW
6. Filter out remaining host sequences using Bowtie2
7. Subsampling to 2 million reads for paired-end data. Subsampling is performed using unique (or deduplicated) reads. NOTE: If a sample no longer has 2 million paired end reads at step 7 it will fail QC.
8. Filter out human sequences using STAR, Bowtie2, and GSNAP
9. Read alignment to NCBI Nucleotide and Protein Database (minimap2 and Diamond)
10. Contig Assembly (SPAdes)
11. Read mapping to contigs for coverage (Bowtie2)
12. Contig Alignment against custom nucleotide and protein databases (BLASTN, BLASTX) (we chose mapping to nucleotide database)
13. Output report of reads and contigs mapped to NCBI taxa as raw reads and read per million mapped reads.

We used the CZID pipeline’s output of reads per million (RPM) for each taxa (species) mapped to the nucleotide database in NCBI. When outputting species level results, all reads mapped to higher orders that could not be resolved at the species level are excluded.

Two input samples failed at this QC level and did not result in taxonomy files - thus our input 697 samples produced 695 taxonomy assignment tables.

## Code framework and implementation

All modeling frameworks were implemented using custom Python pipelines contained in the bio-experiment and bio-sklearn-trainer repositories. Model preprocessing, and hyperparameter configuration are specified through YAML configuration files and training scripts located primarily in GITHUB repository https://github.com/Jtth3Brick/bio-experiment. Scripts for deployment of model training are located in the GITHUB repository https://github.com/Jtth3Brick/bio-sklearn-trainer. This repo contains scripts for data assembly and filtering as well as the model execution scripts responsible for launching parallelized model runs. Configuration files in bio-experiment/experiment_train/config.yaml define the set of preprocessing options, feature selection strategies, model classes, and hyperparameter settings evaluated for each model trained. Hyperparameter exploration was implemented via explicit enumeration of all predefined preprocessing, feature selection, and model parameter combinations specified in the config .yaml file. Each configuration is treated as an independent modeling run and evaluated via 5-fold cross validation.

## Input data, sample selection, and training datasets

All models were trained using species-level metagenomic abundance profiles in reads per million (RPM), obtained from the genomic_sequence_rpm table in the bio.sqlite3 database. To avoid within-subject dependence, a single sequencing run per subject was retained. Sample inclusion and exclusion criteria were applied using metadata stored in the “patients” and “runs” tables with filtering logic implemented in “experiment_train/data_getter.py” as described above in Methods and depicted in Fig.1A. Training datasets were defined as explicit combinations of studies specified in the experiment configuration files “bio-experiment/experiment_train/config.yaml” and denoted as “splits”. These combinations of studies determine which data contribute to the model training datasets.

## Cross-validation strategy

Model performance was evaluated using K-fold cross-validation, with the number of folds specified in the experiment configuration files in “bio-experiment/experiment_train/config.yaml” and recomputed independently for each fold (default = five). Within each fold, samples were partitioned into training and held-out folds. All preprocessing steps, including feature filtering, normalization, and optional feature selection, were fitted exclusively on the training portion of each fold and then applied to the held-out samples.

## Feature filtering and preprocessing

Prior to model fitting, microbial features were filtered using training data to reduce noise and dimensionality. Filtering criteria include A) prevalence-based thresholds, which remove taxa present in only a small fraction of training samples, B) variance-based thresholds, which exclude taxa exhibiting minimal variation across training samples and sometimes C) unsupervised feature selection, which ranks microbial species by importance and retains the top-ranked taxa for downstream model fitting. The specific threshold values used for filtering are defined in the experiment configuration files and applied within each cross-validation fold, without reference to disease labels.

Following filtering, remaining microbial abundance features were normalized on a per-taxon basis using statistics computed from the training samples only. This normalization step places all features on comparable numerical scales prior to modeling and prevents taxa with large numerical ranges from disproportionately influencing model fitting. The same normalization parameters were applied to held-out samples within each fold. All preprocessing and filtering parameters are specified in the configuration files bio-experiment/experiment_train/config.yaml and recomputed independently for each fold.

## Model classes and hyperparameter configuration

7 supervised learning model classes were evaluated. We evaluated seven supervised model classes: L1-regularized logistic regression (lasso), L2-regularized logistic regression (ridge), elastic net logistic regression (ENet), support vector machines (SVC), random forest classifiers (RF), gradient-boosted decision trees (XGB), and feedforward neural networks (NN). Model definitions and hyperparameter ranges are specified in the experiment configuration files bio-experiment/experiment_train/config.yaml and instantiated by the training scripts in the bio-sklearn-trainer repository. For each model class, multiple predefined hyperparameter configurations were evaluated independently within the cross-validation framework. Performance metrics (Cross validation AUC) for the 5 folds were output and saved.

## Model outputs

For each trained model configuration, cross-validation performance metrics and feature importance measures (where able to be extracted) were stored in the structured database tables in bio.sql. These tables include:

train_results: 5-fold CV performance scores for all models run
top_models: top 13 models per class per split (training) selected via 5-fold CV score
feature_importances: features selected for top 5 models per class per split

## Beta-diversity analysis and ordination of gut microbiome profiles

Beta-diversity analyses were performed to evaluate differences in overall gut microbiome composition between individuals with inflammatory bowel disease (IBD) and healthy controls across multiple datasets. All analyses were conducted at the subject level using species-level taxonomic abundance profiles normalized to reads per million (RPM). Samples filtered as noted in section: Filtering Of Metadata Included For Studies. Beta-diversity between samples was quantified using Bray–Curtis dissimilarity computed on the full species-level RPM matrix. Principal coordinates analysis (PCoA) was performed on the resulting distance matrix.

## Feature selection

For all model classes, the top 5 performing models based on CV score were selected for feature extraction. All features included in that model’s prediction were exteacted along with feature weights. From these features the top 80 features per inidvidual model were extracted for downstream analysis. This number of features was selected based on the lowest number of features needed to achieve 95 percent of the feature importance across all model classes evaluated.

## Statistics: PCoA PERMANOVA and PERMDISP

Differences in community composition between IBD and healthy subjects were formally tested using permutational multivariate analysis of variance (PERMANOVA) on the Bray–Curtis distance matrix, with disease status as the grouping variable and 9,999 permutations.

To assess whether observed PERMANOVA results could be driven by differences in within-group dispersion rather than shifts in centroids, permutational analysis of multivariate dispersions (PERMDISP) was also performed using the same number of permutations.

Because datasets differ in population structure and sampling design, PERMANOVA was additionally performed separately within each dataset. For these within-dataset analyses, Bray–Curtis distances were subset to dataset-specific samples and tested using 4,999 permutations.

All beta-diversity analyses were implemented in Python using scikit-bio for Bray–Curtis distance computation, PCoA, PERMANOVA, and PERMDISP, along with pandas, NumPy, matplotlib, and seaborn for data handling and visualization.

## Statistics: DeLong Test of AUC Scores

To assess whether differences in AUC between ensembles were statistically significant, we used the paired DeLong test, which accounts for the correlated nature of predictions on the same evaluation dataset. The test estimates AUC variance and covariance to compute a z-score and p-value. This analysis was conducted using the roc.test() function from the pROC R package, with significance defined as p < 0.05.

## References

1. Bernstein, C. N. & Forbes, J. D. Gut Microbiome in Inflammatory Bowel Disease and Other Chronic Immune-Mediated Inflammatory Diseases. Inflamm. Intest. Dis. 2, 116–123 (2017).

2. Forbes, J. D., Van Domselaar, G. & Bernstein, C. N. The Gut Microbiota in Immune-Mediated Inflammatory Diseases. Front. Microbiol. 7, 1081 (2016).

3. Caron, B., Honap, S. & Peyrin-Biroulet, L. Epidemiology of Inflammatory Bowel Disease across the Ages in the Era of Advanced Therapies. J. Crohns Colitis 18, ii3–ii15 (2024).

4. Fang, X. et al. Gastrointestinal Surgery for Inflammatory Bowel Disease Persistently Lowers Microbiome and Metabolome Diversity. Inflamm. Bowel Dis. 27, 603–616 (2021).

5. Lee, M. & Chang, E. B. Inflammatory Bowel Diseases and the Microbiome: Searching the Crime Scene for Clues. Gastroenterology 160, 524–537 (2021).

6. Franzosa, E. A. et al. Gut microbiome structure and metabolic activity in inflammatory bowel disease. Nat. Microbiol. 4, 293–305 (2019).

7. Liu, J. Z. et al. Association analyses identify 38 susceptibility loci for inflammatory bowel disease and highlight shared genetic risk across populations. Nat. Genet. 47, 979–986 (2015).

8. Clooney, A.G. Ranking microbiome variance in inflammatory bowel disease: A large longitudinal intercontinental study. Gut 70, 499–510 (2021).

9. Bonovas, S. Environmental Risk Factors for Inflammatory Bowel Diseases: An Umbrella Review of Meta-analyses. Gastroenterology 157, 647–659 (2019).

10. Lazar, V. et al. Aspects of Gut Microbiota and Immune System Interactions in Infectious Diseases, Immunopathology, and Cancer. Front. Immunol. 9, (2018).

11. Kaplan, G. G. The global burden of IBD: from 2015 to 2025. Nat. Rev. Gastroenterol. Hepatol. 12, 720–727 (2015).

12. Agrawal, M. & Jess, T. Implications of the changing epidemiology of inflammatory bowel disease in a changing world. United Eur. Gastroenterol. J. 10, 1113–1120 (2022).

13. Burisch, J. Changing epidemiology of immune-mediated inflammatory diseases in immigrants: A systematic review of population-based studies. J. Autoimmun. 105, (2015).

14. Vangay, P. et al. US Immigration Westernizes the Human Gut Microbiome. Cell 175, 962–972.e10 (2018).

15. Deehan, E. C. & Walter, J. The Fiber Gap and the Disappearing Gut Microbiome: Implications for Human Nutrition. Trends Endocrinol. Metab. TEM 27, 239–242 (2016).

16. The Integrative Human Microbiome Project. Nature 569, 641–648 (2019).

17. Serrano Fernandez, V., Seldas Palomino, M., Laredo-Aguilera, J. A., Pozuelo-Carrascosa, D. P. & Carmona-Torres, J. M. High-Fiber Diet and Crohn’s Disease: Systematic Review and Meta-Analysis. Nutrients 15, 3114 (2023).

18. Copeland, J. K. et al. The Impact of Migration on the Gut Metagenome of South Asian Canadians. Gut Microbes 13, 1902705 (2021).

19. Tasson, L., Canova, C., Vettorato, M. G., Savarino, E. & Zanotti, R. Influence of Diet on the Course of Inflammatory Bowel Disease. Dig. Dis. Sci. 62, 2087–2094 (2017).

20. Manichanh, C. et al. Reduced diversity of faecal microbiota in Crohn’s disease revealed by a metagenomic approach. Gut 55, 205–211 (2006).

21. Khorsand, B. et al. Overrepresentation of Enterobacteriaceae and Escherichia coli is the major gut microbiome signature in Crohn’s disease and ulcerative colitis; a comprehensive metagenomic analysis of IBDMDB datasets. Front. Cell. Infect. Microbiol. 12, 1015890 (2022).

22. Lepage, P. et al. Twin Study Indicates Loss of Interaction Between Microbiota and Mucosa of Patients With Ulcerative Colitis. Gastroenterology 141, 227–236 (2011).

23. Halfvarson, J. et al. Dynamics of the human gut microbiome in inflammatory bowel disease. Nat. Microbiol. 2, 17004 (2017).

24. Sharpton, T. J. An introduction to the analysis of shotgun metagenomic data. Front. Plant Sci. 5, (2014).

25. L, M., G, S.-G., Z, X., N, B. & C, M. Intercontinental Gut Microbiome Variances in IBD. Int. J. Mol. Sci. 23, (2022).

26. Dastani, Z. et al. Novel loci for adiponectin levels and their influence on type 2 diabetes and metabolic traits: a multi-ethnic meta-analysis of 45,891 individuals. PLoS Genet. 8, e1002607 (2012).

27. Okada, Y. et al. Genetics of rheumatoid arthritis contributes to biology and drug discovery. Nature 506, 376–381 (2014).

28. Su, Q. et al. Faecal microbiome-based machine learning for multi-class disease diagnosis. Nat. Commun. 13, 6818 (2022).

29. Lloyd-Price, J. et al. Multi-omics of the gut microbial ecosystem in inflammatory bowel diseases. Nature 569, 655–662 (2019).

30. Pasolli, E., Truong, D. T., Malik, F., Waldron, L. & Segata, N. Machine Learning Meta-analysis of Large Metagenomic Datasets: Tools and Biological Insights. PLOS Comput. Biol. 12, e1004977 (2016).

31. Bakir-Gungor, B. et al. Inflammatory bowel disease biomarkers of human gut microbiota selected via different feature selection methods. PeerJ 10, e13205 (2022).

32. Quinn, T. P. et al. A field guide for the compositional analysis of any-omics data. GigaScience 8, giz107 (2019).

33. Zheng, J. et al. Noninvasive, microbiome-based diagnosis of inflammatory bowel disease. Nat. Med. 30, 3555–3567 (2024).

34. Mayorga, L., Serrano-Gómez, G., Xie, Z., Borruel, N. & Manichanh, C. Intercontinental Gut Microbiome Variances in IBD. Int. J. Mol. Sci. 23, 10868 (2022).

35. Smialowski, P., Frishman, D. & Kramer, S. Pitfalls of supervised feature selection. Bioinformatics 26, 440–443 (2010).

36. Kalantar, K. L. et al. IDseq—An open source cloud-based pipeline and analysis service for metagenomic pathogen detection and monitoring. GigaScience 9, giaa111 (2020).

