## Supplement for "Performance of IBD machine learning classifiers varies across microbiome training data independent of geographic diversity"

**Table S1: Patient/sample characteristics for studies post filtering.**
UC= Ulcerative Colitis, CD=Crohn's Disease, IBD=UC+CD patients combined. The dataset name reflects the country where the study was conducted and the last author. Studies where age is not present in the metadata were indicated as >=18 in the primary study's main text.

| Study | Sample Count | M | F | Sex NA | Age NA | Age (Median and IQR) | UC | CD | IBD | Healthy |
|---|---|---|---|---|---|---|---|---|---|---|
| USA_Elinav | 62 | 29 | 32 | 1 | 1 | 38 (28-52) | 8 | 3 | 11 | 51 |
| USA_HMBP | 45 | 20 | 25 | 0 | 0 | 36 (28-50) | 17 | 14 | 31 | 14 |
| USA_PRISM | 212 | 35 | 40 | 137 | 0 | 35 (28-55) | 86 | 92 | 178 | 34 |
| Ireland_Hill | 61 | 23 | 30 | 8 | 8 | 36 (29-48) | 12 | 5 | 17 | 44 |
| Germany_Elinav | 182 | 77 | 105 | 0 | 0 | 55 (43-63) | 59 | 0 | 59 | 123 |
| Israel_Elinav | 64 | 26 | 27 | 11 | 21 | 37 (26-49) | 13 | 23 | 36 | 28 |
| China_Jia | 69 | 57 | 12 | 0 | 0 | 25 (23-31) | 0 | 42 | 42 | 27 |

**Table S2:Within-cohort PERMANOVA results (Healthy vs IBD) on filtered shotgun metagenomic data.**
Within-study PERMANOVA analyses were performed for each data set (groupings = Healthy and IBD = Ulcerative colitis + Crohn's disease samples) using Bray–Curtis dissimilarities and 4,999 permutations. Input samples in read per million normalized counts tables filtered and processed as per Fig 1A.

| Study | Samples | Healthy | IBD | pseudo-F | p-value | $R^2$ |
|---|---|---|---|---|---|---|
| USA_Elinav | 62 | 51 | 11 | 2.46 | $5.4x10^{-3}$ | 0.039 |
| USA_HMBP | 45 | 14 | 31 | 0.92 | $5.1x10^{-1}$ | 0.021 |
| USA_PRISM | 212 | 34 | 178 | 5.71 | $2x10^{-4}$ | 0.026 |
| Ireland_Hill | 61 | 44 | 17 | 3.52 | $1x10^{-3}$ | 0.056 |
| Germany_Elinav | 182 | 123 | 59 | 10.84 | $2x10^{-4}$ | 0.057 |
| Israel_Elinav | 64 | 28 | 36 | 4.08 | $2x10^{-4}$ | 0.062 |
| China_Jia | 69 | 27 | 42 | 7.24 | $2x10^{-4}$ | 0.097 |

**Table S3: Distribution of models trained per training dataset, by model class.**
For each of the 5 training datasets all 7 model classes were trained. For each class we show the average and sd of the numbers of models run for that class across all 5 training datasets. Differences in number of model classes run between model classes reflect the size of the hyperparameter search per model class (see methods).

| Model class | Average model count across all training datasets (mean ± SD) |
|---|---|
| RF | 1,469 ± 3 |
| XGB | 6,595 ± 103 |
| lasso | 2,280 ± 0 |
| ridge | 2,279 ± 2 |
| ENet | 8,177 ± 66 |
| neural networks | 24,178 ± 27 |
| SVC | 4,559 ± 1 |

**Table S4: SRA accession numbers and citations for studies included in modeling and evaluation.** Sequencing read archive ascension numbers and study citations for accessing raw metagenomic samples and metadat for patients in each study. Where multiple ascession numbers are used patients are from different studies but the same medical center/IBD patient research group.

| Study Name | SRA Number | Reference |
|---|---|---|
| USA_Elinav | PRJEB50555 | Frederici et al., 2022 |
| USA_HMBP | PRJNA400072 | Khorsand, Babak et al, 2022 |
| USA_PRISM | PRJNA400072<br>PRJNA384246 | Ananthakrishnan et al., 2017<br>Franzosa et al. 2019 |
| Ireland_Hill | PRJNA813736 | Stockdale et al., 2022 |
| Germany_Elinav | PRJEB50555 | Frederici et al., 2022 |
| Israel_Elinav | PRJEB50555 | Frederici et al., 2022 |
| China_Jia | PRJEB15371 | He et al., 2017<br>Mayorga et al., 2022 |

**Fig S1: dataset-specific separation of IBD and healthy subjects in Bray–Curtis PCoA space**
Principal coordinates analysis (PCoA) of Bray–Curtis dissimilarities computed from species-level, reads-per-million–normalized gut microbiome profiles. Each panel shows samples from a single dataset projected into a shared global PCoA space, with points colored by disease status (IBD vs. healthy). The same coordinate axes and scaling are used across all panels to enable direct visual comparison of cohort-specific separation patterns. Axis labels indicate the percentage of variance explained by each principal coordinate. IBD = UC + CD combined.

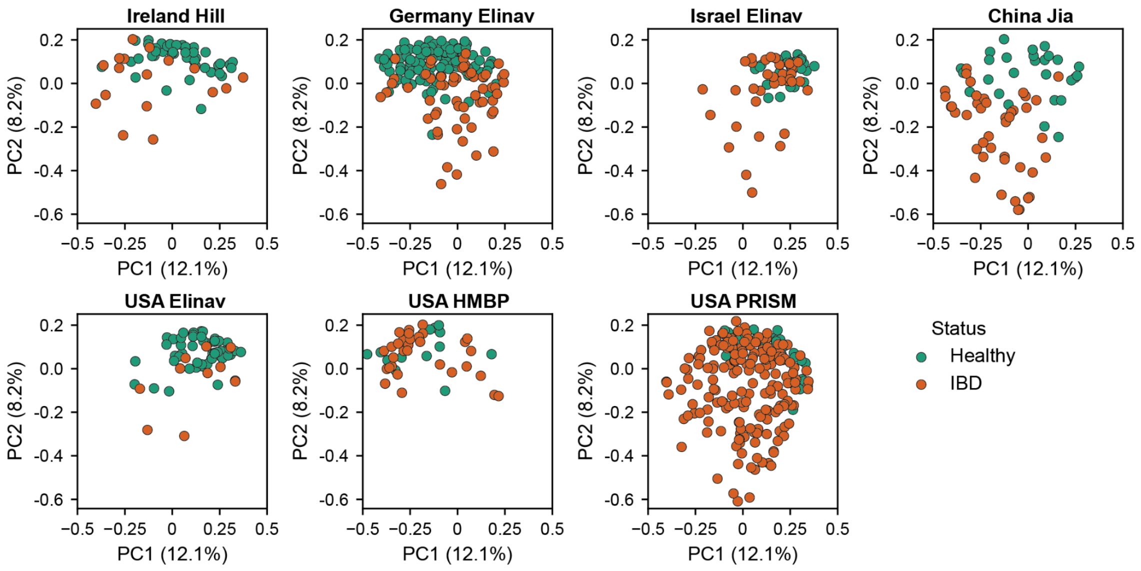


**Fig S2. Performance across international evaluation studies separating out strong model classes (extreme gradient boosting and random forest).**

For each evaluation study, AUC values were computed across all applicable training datasets for single model class ensembles (13 top models). Boxplots are separated by high performing single model-ensemble classes (XGB and RF) and lower performing model classes (ridge, lasso, neural net, ENet and SVC = "all other models"). Boxes indicate the median and interquartile range (IQR), with whiskers extending to 1.5 IQR. Outlier points placed on x-axis.

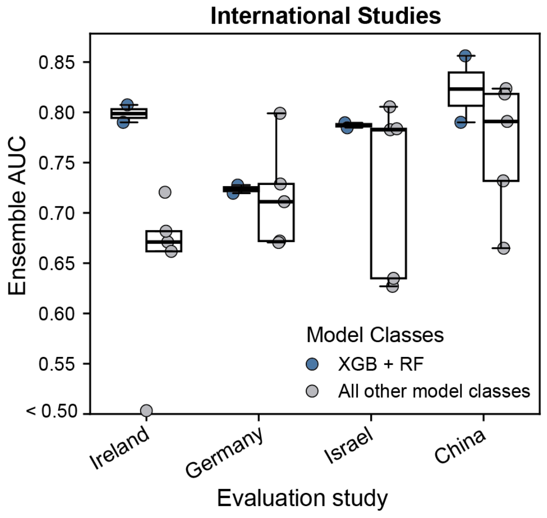


**Fig S3 Performance across USA evaluation studies separating out strong model classes (extreme gradient boosting and random forest).**

For each evaluation study, AUC values were computed across all applicable training datasets for single model class ensembles (13 top models). Boxplots are separated by high performing single model-ensemble classes (XGB and RF) and lower performing model classes (ridge, lasso, neural net, ENet and SVC = "all other models'). Boxes indicate the median and interquartile range (IQR), with whiskers extending to 1.5 IQR.

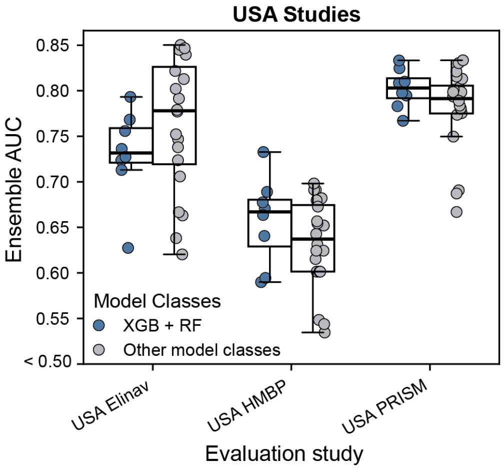


**Figure S4 Performance on USA-datasets for all models, split by international training datasets 1-4.**

For each single model class ensemble (top 13 models per class), AUC scores (points) for each USA study are shown. Data plotted separately per international training dataset (Int_1 to Int_4). Boxes indicate the median and interquartile range (IQR), with whiskers extending to the full range.

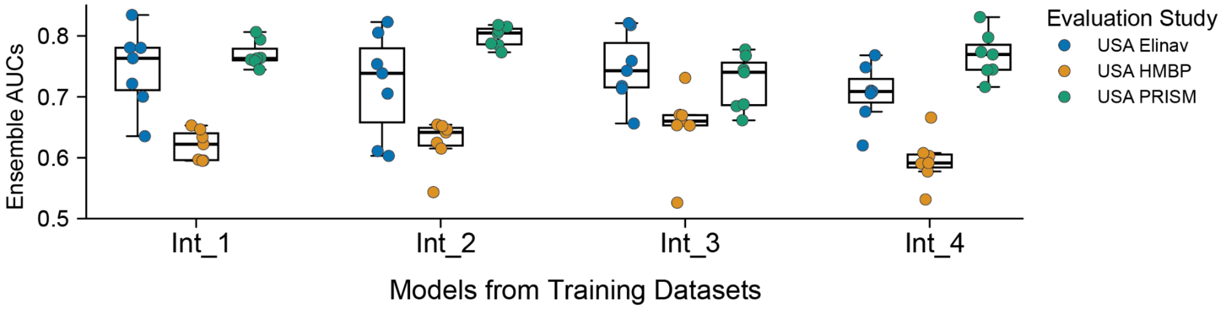


**Figure S5: Median cumulative feature-importance curves by model class, with 95% contribution thresholds**

Median cumulative feature-importance curves for each model class. Medians calculated from 25 models (top 5 per training dataset across all 5 datasets). For each model class, curves were pooled across configurations and summarized by their median. Features were ranked by absolute importance within each model configuration, and cumulative importance was normalized to sum to one. The x-axis shows the fraction of ranked features included, and the y-axis shows the cumulative fraction of total importance captured. Dots indicate where the median curve reaches 95% of cumulative importance; legend labels report the estimated number of features required to reach this threshold per model class.

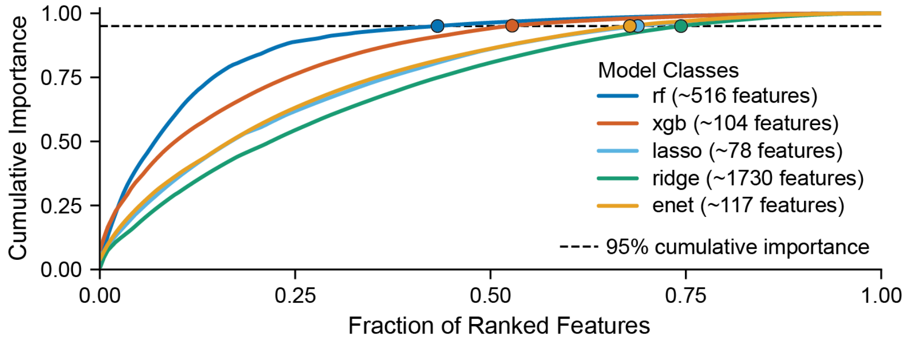


**Figure S6: Unique species selected per training dataset - by model class**

Number of unique microbial species selected in each training dataset after applying the per-model top-80 feature (species) filter to all 5 top models per model class. For all 5 top models per model class, the set of unique species per model class was retained. Bars show totals unique species per training dataset (USA; Int_1–Int_4) stacked by the model class.

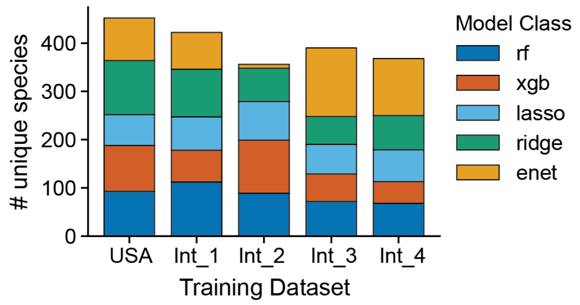


**Figure S7 : Bar plot showing the number of microbial taxa shared as top features across training datasets.**

For each training dataset, the considered microbial taxa (features) are the intersection of the top 80 important features across all model classes. Species are counted and binned by how many training datasets share that species (1,2,3,4 or 5).

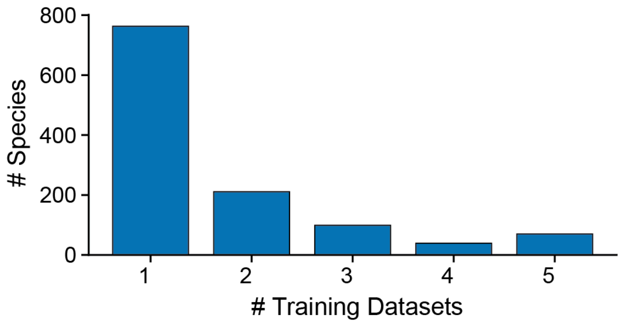


**Figure S8: Principal Coordinates Analysis (PCoA) of study-associated microbiome structure across all filtered samples.**

PCoA based on Bray-Curtis dissimilarity was performed across all filtered samples included in the study (see Fig. 1A). Each point represents the gut microbiome of a single individual and is colored according to study identity.

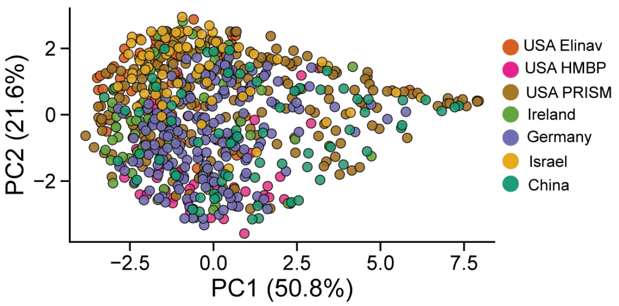
PC2 (21.6%)
2
0
−2
−2.5
0.0
2.5
5.0
7.5
PC1 (50.8%)
USA Elinav
USA HMBP
USA PRISM
Ireland
Germany
Israel
China
